# Microfluidics-Enabled Simultaneous Imaging of Neural Activity and Behavior in Chemically Stimulated, Head-Fixed *C. elegans*

**DOI:** 10.1101/2025.11.03.686171

**Authors:** Hyun Jee Lee, Julia Vallier, Hang Lu

## Abstract

Understanding how the brain processes sensory information and produces appropriate behavior is a fundamental question in neuroscience. In this study, we developed a novel microfluidic device that allows for simultaneous observation of neural activity and behavior in *Caenorhabditis elegans* (*C. elegans*) during chemosensory stimulation. Traditional methods often involve trade-offs between high-resolution neuronal imaging, behavioral recording, and the ability to apply chemical stimulation. Our innovative design overcomes these limitations by immobilizing the worm’s head, stabilizing neuronal imaging, while allowing the posterior portion of the body to move freely, enabling the study of naturalistic behaviors during chemical stimulation. We applied this device to investigate how *C. elegans* responds to both attractive and aversive chemical cues. By correlating neural activity with observed behavior, we identified neurons and whole-brain dynamics associated with specific movements. Our results demonstrate that providing the worm with greater freedom of movement results in more naturalistic neuronal and behavioral responses to stimuli, compared to fully immobilized setups. This new tool offers a powerful approach for studying how sensory information is processed in the *C. elegans* nervous system to generate behavior, with potential applications in other model organisms. Its versatility and ease of operation make this device broadly applicable for studying how neural circuits drive behavior and decision-making in complex environments.

## Introduction

A central goal in neuroscience is to explain how neural dynamics give rise to behavior^[1,2]^. This relationship forms the foundation for comprehending fundamental brain functions such as perception, decision-making, movement, and learning ^[3–5]^. Behaviors are not the result of isolated neurons acting independently but are instead generated by networks of neurons working together in circuits, dynamically processing sensory information and coordinating motor outputs ^[6–8]^. To fully understand these processes, it is important to measure the activity of large populations of neurons in moving animals during naturalistic behaviors like sensing, motor execution, and decision making ^[9–11]^.

*Caenorhabditis elegans* (*C. elegans*) is an ideal model organism for understanding how genes and environment shape behavior via neural circuit activities. With a compact nervous system comprising only 302 neurons, *C. elegans* allows for multi-neuronal calcium imaging at cellular resolution, an approach that is currently infeasible in more complex mammalian brains ^[12–14]^. Despite its simplicity, *C. elegans* shares significant homology with humans, and many of the cellular and molecular mechanisms governing its nervous system, such as synaptic transmission, are highly conserved ^[15–19]^. Moreover, *C. elegans* exhibits a wide range of behaviors in response to diverse external stimuli, including chemical, mechanical, and thermal cues, making it a valuable model for investigating how sensory-induced neural dynamics drive behavior ^[20–22]^.

In *C. elegans*, chemosensation, encompassing both olfactory and gustatory pathways, is the primary modality through which the organism perceives and responds to its environment, playing a key role in behaviors such as foraging, threat avoidance, and social interactions ^[23,24]^. Chemical stimulation has long been used to study behaviors and neuronal activity related to chemosensation, chemotaxis, habituation, and learning ^[25–27]^. Microfluidic techniques are commonly employed for this purpose, as they enable precise manipulation of both the organism and its environment, allowing for the controlled delivery of chemical stimuli with high temporal and compositional accuracy ^[20,21,28]^. The most commonly used microfluidic configuration involves immobilizing the worm in a narrow microchannel, with its head positioned at the site where the chemical stimulus is applied ^[20,21]^. This physical restraint of the body stabilizes head motion, offering several advantages: it allows high-magnification imaging of neurons in the head region without losing them from the field of view, minimizes motion artifacts in neural recordings for high-fidelity data acquisition, and ensures the consistent delivery of chemical stimuli to the sensory organs in the nose. However, this design also presents limitations. While immobilization facilitates stable imaging, it constrains the worm’s body, preventing observation of its behavioral responses to stimuli, which are critical for understanding how sensory inputs translate into motor outputs. Without the ability to monitor behavior, it becomes difficult to correlate neural activity with the worm’s perception and decision-making processes. Moreover, physical restraint may affect the worm’s response via proprioception, potentially altering neural responses in the underlying neural circuits ^[13,29]^. To address this, separate behavioral assays are often conducted on a different population of worms in parallel experiments ^[30,31]^. However, this approach only allows for indirect comparisons between averaged neural activity and averaged behavioral responses, without establishing a direct link between individual neuronal activity and behavior within the same worm. Additionally, the behaviors observed in separate assays, often conducted under different experimental conditions (e.g., on plates rather than in microfluidic channels), may not accurately reflect those that occur during on-chip chemical stimulation, further complicating the interpretation of results.

Other existing platforms also fall short of enabling simultaneous multi-neuronal imaging and behavior observation during chemical stimulation. Micropillar array systems, which allow *C. elegans* to freely roam in a chemically controlled liquid environment, have been used to correlate behavior with neural activity in freely moving worms during chemical stimulation ^[28,32,33]^. However, due to the wide-field imaging required to capture the entire array area, these systems suffer from low spatial resolution, limiting their ability to image large populations of neurons. This is a critical drawback, as behavior is driven by large networks of neurons, making the imaging of many neurons necessary to understand the neural basis of behavior. Consequently, micropillar array systems have been primarily used for single neuron or very sparsely labeled neuron imaging. On the other end of the spectrum, specialized microscopy platforms have been developed that are capable of high-resolution functional imaging while automatically tracking freely moving *C. elegans* ^[34,35]^. These systems use high-resolution volumetric imaging to capture the neurons in the head, while separate objectives and cameras are used to image the worm’s movement at lower resolution. Real-time feedback loops are employed to track the head’s position in three dimensions (x, y, z), keeping it centered in the field of view. However, these platforms are not designed to accommodate chemical stimulation within microfluidic devices. The systems cannot handle the thickness of PDMS in microfluidic set-ups or the rapid movements of worms in liquid environments, making it difficult to track neural activity without motion artifacts in real time. Even if this limitation were overcome, implementing rapid changes in liquid composition around the entire body of a freely moving worm would require high flow rates that can stress the animal and alter its natural behavior. Additionally, these platforms involve complex and less accessible instrumentation, as well as substantial image processing post-acquisition to align high-resolution images and track neurons ^[36]^. Therefore, there is a need for a new approach to integrate all three critical functions: chemical stimulation, multi-neuronal imaging, and behavioral recording in moving *C. elegans*.

In this paper, we present a novel microfluidic device that enables simultaneous imaging of multi-neuronal, including whole-brain, activity and behavior during chemical stimulation in *C. elegans*. Our approach involves selectively immobilizing only the head of the worm while allowing the posterior part of the body to move freely, enabling the observation and interpretation of its behavior in real-time. This technique is analogous to head-fixation methods used in other model organisms, such as *Drosophila*, zebrafish, and mice, where the head is immobilized while other parts of the body are free to move ^[37–39]^. Head fixation in these models has proven to be highly effective in facilitating high-resolution neural imaging without the motion artifacts typically encountered in freely moving animals. Developing such a system for *C. elegans* presents unique challenges due to its physical characteristics: it is small (1 mm in length), soft, cylindrically shaped, and moves quickly (up to 4.5 body lengths per second) ^[40]^. Previously, we developed a head-immobilization method based on localized hydrogel polymerization around the worm’s head ^[41]^. While this technique effectively immobilized the head, the hydrogel chemistry limited the range of chemical stimuli that could be used and was not ideal for long-term imaging due to osmolarity-induced stress on the worm. The new device we present in this work overcomes these limitations by using a purely hydrodynamics-driven approach, eliminating the need for polymerization chemistry. We demonstrate that the head fixation is robust across various conditions and can be applied in a wide range of experimental applications. We observed reliable behavioral and neuronal responses to both attractive and aversive chemical stimuli. Additionally, we show that the device is compatible with standard epifluorescence microscopy, standard confocal microscopy as well as custom-built light-sheet microscopy, making it accessible to a broad range of research laboratories. Notably, our method improves neuronal responsiveness compared to conventional microfluidic devices that fully restrain the worm’s body, likely because the freedom of tail movement allows for more natural sensory and behavioral responses. This system thus addresses a significant gap in current methods by enabling simultaneous chemical stimulation, high-resolution multi-neuronal imaging, and behavioral recording in *C. elegans*, providing a powerful tool for studying the neural mechanisms underlying behavior.

## Results

### Observing Neuronal Activity and Behavior During Chemosensory Stimulation

With the goal of developing a *C. elegans* version of the head-fixation method, we engineered a microfluidic device that enables simultaneous imaging of neuronal activity and behavioral recording during chemical stimulation (Figure 1A). Head fixation is crucial for obtaining high-fidelity neuronal signals, as it minimizes motion artifacts that typically compromise imaging quality in moving animals. This stabilization allows us to image multi-neuronal activities at single-cell resolution, significantly enhancing the accuracy and reliability of data collection (Figure 1B). Moreover, the stability of the head allows the precise and localized delivery of chemical stimuli to the anterior region, where chemosensory neurons are predominantly concentrated. The device’s chemical stimulus delivery system is similar to that of conventional designs, positioning the worm’s head towards the T-junction where the stimulus is applied ^[21]^. However, our device incorporates a shorter narrow channel that fits only the anterior portion of the worm, while leaving the tail free to move in a wider chamber. The head-fixation mechanism relies solely on fluid mechanics, eliminating the need for active control using additional software or hardware components beyond those used in conventional devices. This design ensures that the device operation remains robust, straightforward, and accessible for a wide range of experimental applications. While the worm remains stationary overall, the direction of wave propagation in the tail provides key behavioral information: forward locomotion is indicated by wave propagation from head to tail, and backward locomotion is signaled by wave propagation from tail to head (Figure 1C). Additionally, other behavioral responses, such as curling and tail pushing against the chamber walls, are observed and can provide further context for interpreting neuronal activity. This ability to observe and interpret behavior, which reveals the animal’s perception of the chemical stimulus, was not achievable with conventional devices that fully immobilize the worm.

**Figure 1:**
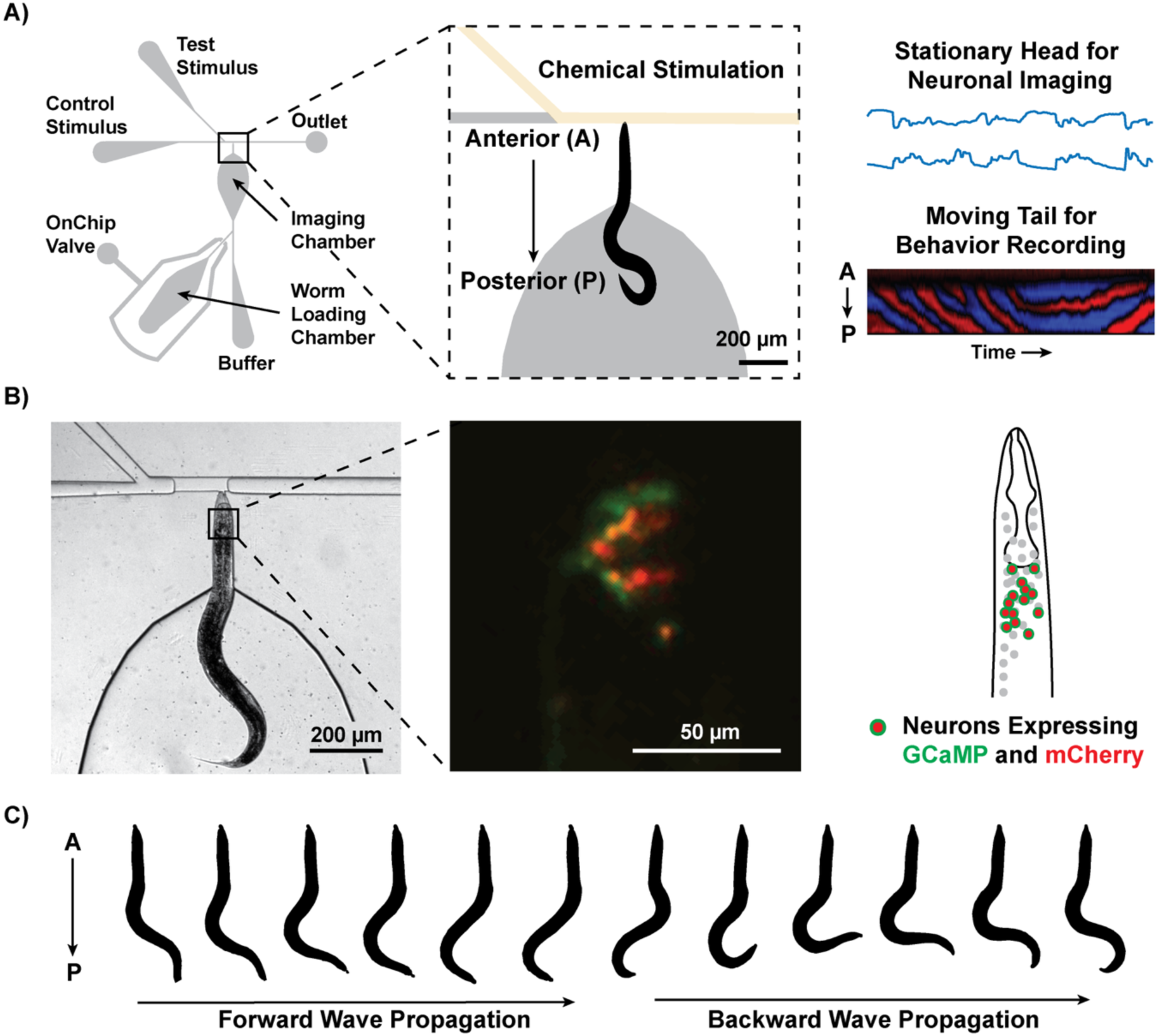
Device design. A) The device is designed to enable simultaneous recording of neural activity and behavior during chemical stimulation. The anterior portion of *C. elegans* is immobilized in a narrow channel, while the posterior portion remains free to move in a wider section of the imaging chamber. Chemical stimuli are precisely delivered to the anterior end via microfluidic actuation. B) Example brightfield image of a head-fixed worm on chip and a fluorescent image of the head region containing multiple neurons expressing GCaMP and mCherry. C) The worm exhibits both forward and backward movements, which are inferred from the direction of wave propagation along the body.

### Fluid Mechanical Characterization of Device Operation

The key to localized immobilization in our system relies on the Venturi effect, a fluid mechanical phenomenon that describes the pressure drop occurring when fluid flows through a constriction ^[42]^. As fluid flows from the wide imaging chamber into the narrow channel, this pressure drop creates a localized vacuum that securely holds the worm’s head in place, preventing escape. This process arises automatically from the device geometry, ensuring simple and robust operation. To ensure consistent flow direction, a small amount of pressure is applied to the imaging chamber. It is important to note that a small amount of fluid continues to flow around the head due to the rectangular shape of the channel, even when the worm is securely positioned. The width and shape of the imaging chamber were specifically designed to accentuate the pressure drop: it is as wide as possible to enhance the pressure differential while directing the flow into the narrow channel. Additionally, the constricted end of the channel features a three-step vertical taper to further increase the constriction effect and accommodate the worm’s head shape.

To validate the presence and functionality of the Venturi effect, we simulated the pressure profile within the device using COMSOL (Figure 2A). A three-dimensional model of the T-junction region, incorporating the three-step vertical taper, was developed to replicate the device’s fluid dynamics assuming incompressible laminar flow. The simulation confirmed a significant pressure drop along the narrow channel, with the lowest pressure observed at the T-junction, consistent with the principles of the Venturi effect. This pressure differential is key to securely immobilizing the worm’s head. To empirically verify the pressure drop, we employed particle velocimetry to track the movement of beads flowing within the device (Figure 2B). The flow velocity field throughout the device was calculated from the trajectories of micro-sized beads suspended in buffer, captured using a high-speed camera. The results revealed a pronounced velocity gradient along the narrow channel, further corroborating the presence of the Venturi effect (Figure 2C). In contrast, the flow velocity within the wider portion of the imaging chamber was notably low, ensuring minimal disruption to the tail’s natural movement. We observed no behavioral effects on tail movement due to this minor flow, demonstrating that the device effectively stabilizes the worm’s head without compromising behavioral fidelity.

**Figure 2:**
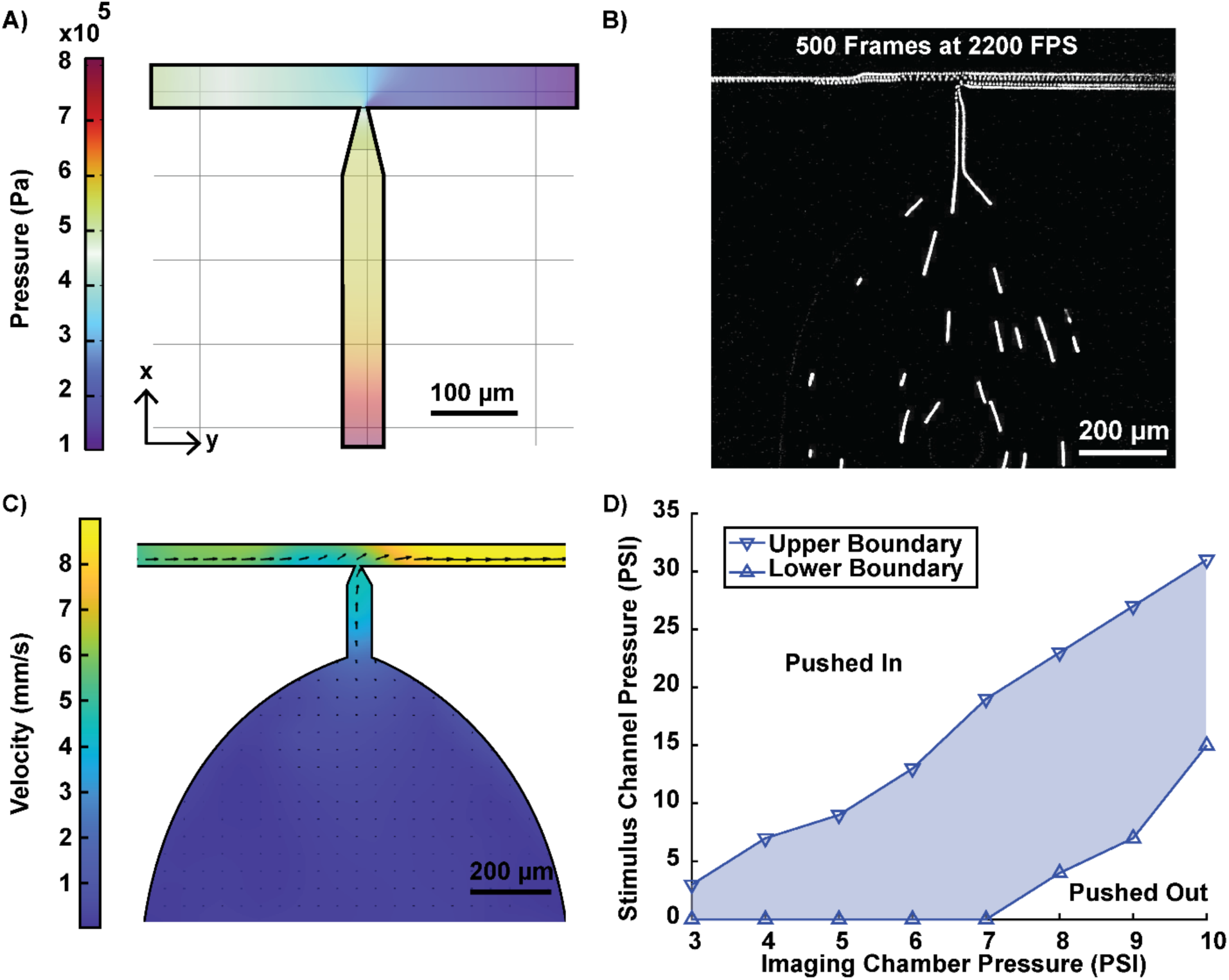
Characterization of the Venturi effect and operating conditions. A) Simulation of the pressure decrease due to flow through a constriction, illustrating the Venturi effect. B) Trajectories of microbeads inside the device in 0.2 seconds visualized by max projection of consecutive frames. C) Experimental velocity profile within the device, calculated using particle image velocimetry. D) Operating pressure range of the device. The blue shaded area represents the pressure balance between the imaging chamber and stimulus channel that ensures proper worm positioning. Pressure combinations above the upper boundary line cause the worm to be pushed into the imaging chamber, while pressure combinations below the lower boundary push the worm toward the outlet.

Maintaining the Venturi effect, and thereby stable head fixation, relies on achieving an appropriate balance between the pressures in the imaging chamber and the stimulus channel. To identify the optimal operating conditions, we systematically tested various pressure combinations to ensure reliable head immobilization. Remarkably, head fixation was maintained over a broad range of pressures (Figure 2D). For example, when the imaging chamber pressure was set to 8 psi, the stimulus channel pressure could range from 4 to 23 psi. Pressures below this range caused the worm to be pushed out of the narrow channel, while pressures above it forced the worm back into the wider imaging chamber. In most experimental setups, an imaging chamber pressure of 8–9 psi and a stimulus channel pressure of 4–10 psi provided consistent fixation. These operating pressures fall well within the range commonly used in microfluidic systems. The wide operational range of the device indicates that it is not only highly adaptable but also exceptionally robust. In other words, minor fluctuations in pressure, which can occur due to environmental or equipment variability, do not disrupt head fixation or compromise the worm’s immobilization. Furthermore, the timing and composition of chemical stimuli were preserved across these conditions (Figure S1). This robustness ensures precise control over chemical delivery, even in varying experimental conditions. Overall, the ability to maintain stable head fixation across a reasonable and forgiving range of pressures makes this device a practical and versatile tool for diverse microfluidic applications, from neuronal imaging to behavioral analysis. The reliability of fixation under these conditions reduces experimental variability, streamlining data collection and enhancing reproducibility across experimental setups.

### Exploring Different Modes of Device Operation

The robustness of the device operation enables flexibility in design and supports various modes of operation. By tuning the length of the narrow channel, we sought to investigate how different degrees of body immobilization influence the worm’s movement patterns and kinematics. Channels were fabricated to immobilize approximately 30%, 40%, and 60% of the body length of a day-1 adult hermaphrodite *C. elegans* (Figure 3A). These proportions were carefully chosen to balance two factors: providing enough restraint to stabilize anterior neurons for high-fidelity imaging, while leaving the posterior region sufficiently free to exhibit natural movement. The shorter channels (30% and 40%) primarily immobilized the anterior region, which houses the majority of sensory and interneurons, while leaving most of the posterior, including the tail, free to move. In contrast, the longer channel (60%) extended the restraint toward the mid-body, restricting regions associated with egg-laying and movement generation, leaving only the extreme posterior free.

**Figure 3:**
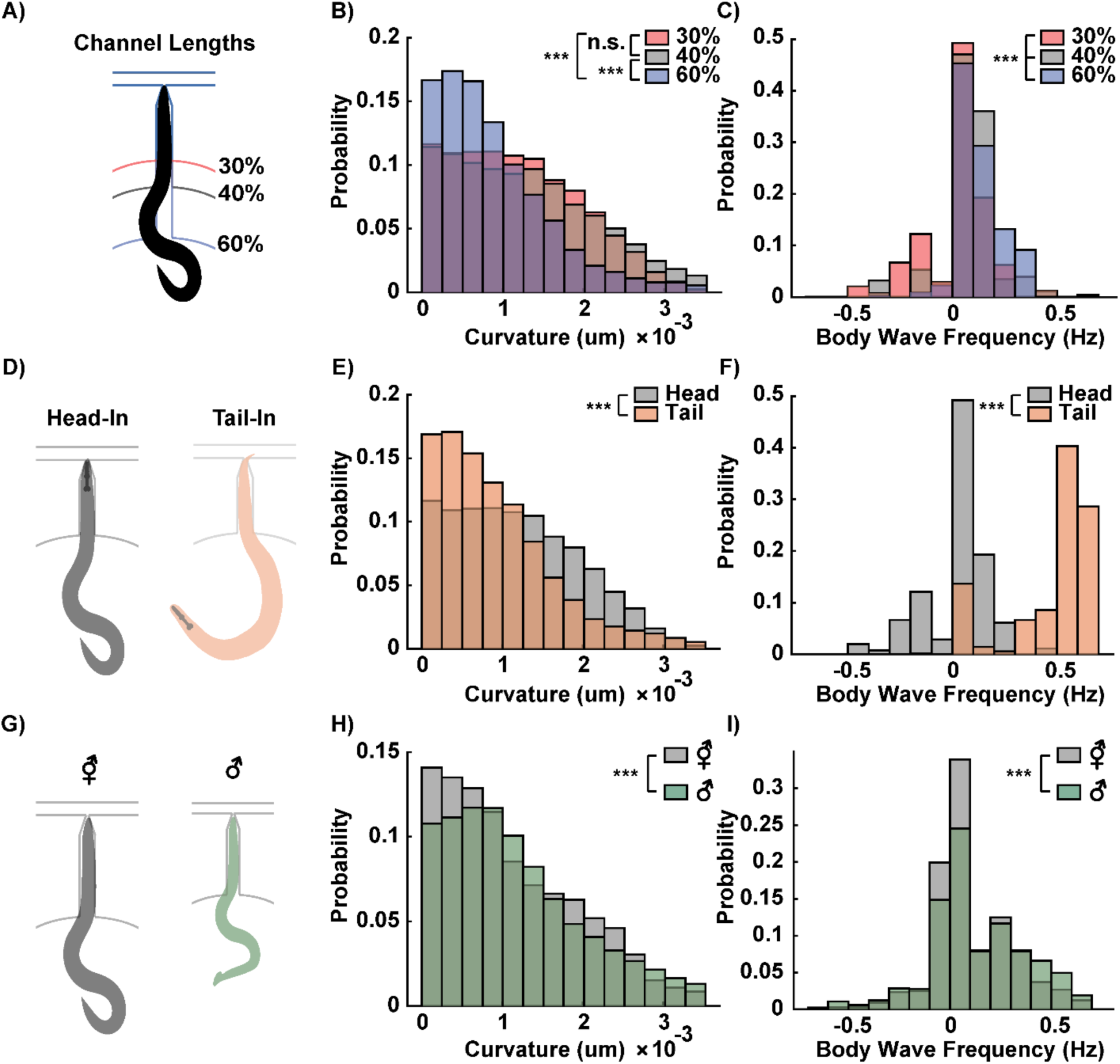
Modes of operation and resulting body curvature and wave frequency distributions. A) The length of the narrow channel is adjusted to immobilize approximately 30%, 40%, or 60% of the worm’s body. B) Histograms of body curvature in the freely moving portion of the worm’s body. While 30% and 40% immobilization show no significant difference in curvature, 60% immobilization results in significantly lower average curvature in the posterior region. C) Histograms of body wave frequency. Worms with 30% immobilization exhibit a significantly lower mean wave frequency compared to those with 40% immobilization. D) The device can also immobilize the tail end of the worm in the narrow channel, allowing for tail neuron stimulation and imaging. E, F) Curvature and body wave frequency distributions differ significantly between head-fixed and tail-fixed animals in the 30% immobilization configuration. G) The overall dimensions of the device are adjusted to accommodate animals of different sizes, such as hermaphrodite and male worms of the same age (day 3). H, I) Smaller and thinner male worms exhibit higher average curvature and body wave frequency compared to hermaphrodites of the same age. Two sample t-tests were performed for statistical analysis. The asterisk symbols denote a significance level as follows: P < 0.05 = *, P < 0.01 = **, P< 0.001= ***.

Differences in body curvature and wave propagation across channel designs provided insights into the mechanical, proprioceptive, and neural constraints imposed by partial physical restraint. The distribution of curvature in the unrestrained body segments showed a significantly lower curvature in worms loaded into the 60% channel compared to the 30% and 40% channels (Figure 3B). No significant difference in curvature was observed between the 30% and 40% channels. Wave propagation also varied with channel length. Increased immobilization reduced the likelihood of backward wave propagation, characterized by negative body wave frequency (Figure 3C). Backward wave propagation was most frequently observed in the 30% channel, where the tail retained the greatest degree of freedom.

The observed differences in curvature and wave frequency across channel lengths may stem from several factors, including biomechanics, proprioception, and neural control. First, the increased restraint in the 60% channel physically restricts posterior movement, limiting the ability to generate high curvature and coherent backward waves. Second, proprioceptive feedback could also play a role, as the extent of body contact with the microchannel may alter sensory input and motor output. For example, the anterior-focused mechanical stimulation in the 30% channel may induce escape responses, manifesting as increased backward movement ^[43,44]^. Finally, the reduction in backward wave propagation in the 60% channel could result from decoupling or modulation of distributed rhythm generators, which are critical for coordinating coherent wave propagation along the body ^[45,46]^.

These findings also suggest that different channel lengths can be tailored to meet specific experimental needs: the 30% channel can be used to study backward wave propagation in greater focus, while the 60% channel can be used to suppress curling motions. It is important to note that, in some cases, worms in the 30% channel were able to push their heads out of the narrow channel by exerting forces with their tails against the channel walls. However, the worm could not fully escape, as the Venturi effect would pull the tail into the channel as soon as the head exited, switching from head immobilization to tail immobilization. For most neural imaging experiments, we used the 40% channel, as it provided a balance between minimizing the restriction of body curvature while ensuring stable head immobilization. This channel length allowed us to observe a range of behaviors, including forward, backward, and coiling movements. The tunability of immobilization in the device not only provides flexibility for neural imaging but also holds potential for studying the interplay between biomechanics, proprioception, and neural circuits in locomotion.

Imaging the tail region of the worm offers additional opportunities to probe neural circuits involved in sensory processing and behavior. While the majority of *C. elegans* neurons (∼180) are located in the head, approximately 45 neurons are in the tail, including chemosensory neurons and others that play roles in escape and reproductive behaviors ^[47]^. For example, PHA and PHB chemosensory neurons detect noxious chemicals, while the PCA and PCB neurons are involved in mating behaviors ^[48,49]^. Our device can also immobilize the tail region within the narrow channel while leaving the anterior portion of the worm free to move (Figure 3D), enabling tail-focused imaging or localized chemical stimulation. The direction in which the worm enters the narrow channel (whether head-fixed or tail-fixed) is initially random but can be controlled by toggling the Venturi suction: temporarily reversing the flow allows the animal to turn, after which suction can be reactivated to secure the desired head-fixed or tail-fixed orientation.

Our findings show statistically significant differences in the curvature of the freely moving body segments depending on whether the head or tail was immobilized (Figure 3E). Specifically, curvatures in the freely moving tails of head-fixed worms were higher on average than the curvatures in the freely moving heads of tail-fixed worms, suggesting that the posterior region of the worm is more flexible and capable of sharper curls than the anterior portion. This difference in flexibility may be attributed to biomechanical differences between the regions, such as the rigidity imparted by the pharynx in the anterior or the more densely innervated motor neuron network in the head region ^[15]^. Interestingly, tail to head wave propagation was not observed in tail-fixed worms (Figure 3F). Instead, these worms exhibited high-frequency forward movements, as indicated by high positive body wave frequency values. This observation may be reflecting forward escape behavior of worms when they are stimulated in the tail ^[50]^. It may also suggest that immobilizing the tail inhibits the normal neural and muscular coordination required for backward locomotion.

Beyond controlling head or tail immobilization, the device can be readily scaled to accommodate worms of different sizes, enabling comparisons across developmental stages and sexes. For example, the device originally designed for day 1 hermaphrodite worms was scaled by 120% to fit larger day 3 hermaphrodite worms (Figure 3G). This scalability allowed us to compare the behavior of day 3 hermaphrodite and male worms under the same experimental conditions, with both groups experiencing 40% body immobilization. Male worms, which are shorter, thinner, and lacking eggs in the mid-body, exhibited higher curvature in their tails compared to their hermaphrodite counterparts (Figure 3H). The presence of eggs, which are mechanically more rigid, can reduce the body’s flexibility, limiting its ability to exhibit high curvature. Additionally, males demonstrated higher average body wave frequencies (Figure 3I), possibly due to their reduced mechanical constraints, as they are thinner and do not carry eggs. These findings may also be related to the known differences in sex-specific behaviors and biomechanics in *C. elegans*, such as increased curling motions and exploratory behavior in males driven by sex-specific circuits ^[51,52]^. The flexibility of the device design provides a platform for addressing a wide range of experimental questions, such as how sex and developmental stage influence the neural and biomechanical coordination of movement. This adaptability extends the utility of the device beyond calcium imaging and behavioral recording during chemical stimulation, offering potential applications in studies of developmental biology, sexual dimorphism, and the integration of neural and mechanical systems in behavior.

### Observing Neuronal Activity and Behavior During Chemosensory Stimulation

Imaging a *C. elegans* in our device allows us to link neuronal activity to the worm’s concurrent behavior, providing information comparable to that obtained from freely moving animals. This direct correspondence between neural dynamics and motor output offers a powerful framework for understanding how neural circuits drive behavior. To represent the worm’s behavior over time, the body shape is segmented from the video recording, and the centerline is extracted. The centerline is then divided into segments, and the curvature of each segment is mapped to generate a kymograph (Figure 4A). This curvature kymograph provides a qualitative visualization of wave propagation direction, wave frequency, and curvature magnitude. When exposed to two cycles of an attractive food odor stimulus, the worm exhibited forward locomotion during the stimulus, indicating an attempt to approach the favorable odor. Upon removal of the food odor, the worm switched to backward locomotion, a behavior consistent with an aversive response to the absence of the positive stimulus. It is important to note that behavior is not strictly deterministic and often involves delays in response. These observations suggest that the worm’s behavior dynamically reflects its perception of the stimulus, with forward movement representing an attraction to the odor and reversals indicating dissatisfaction with its removal.

**Figure 4:**
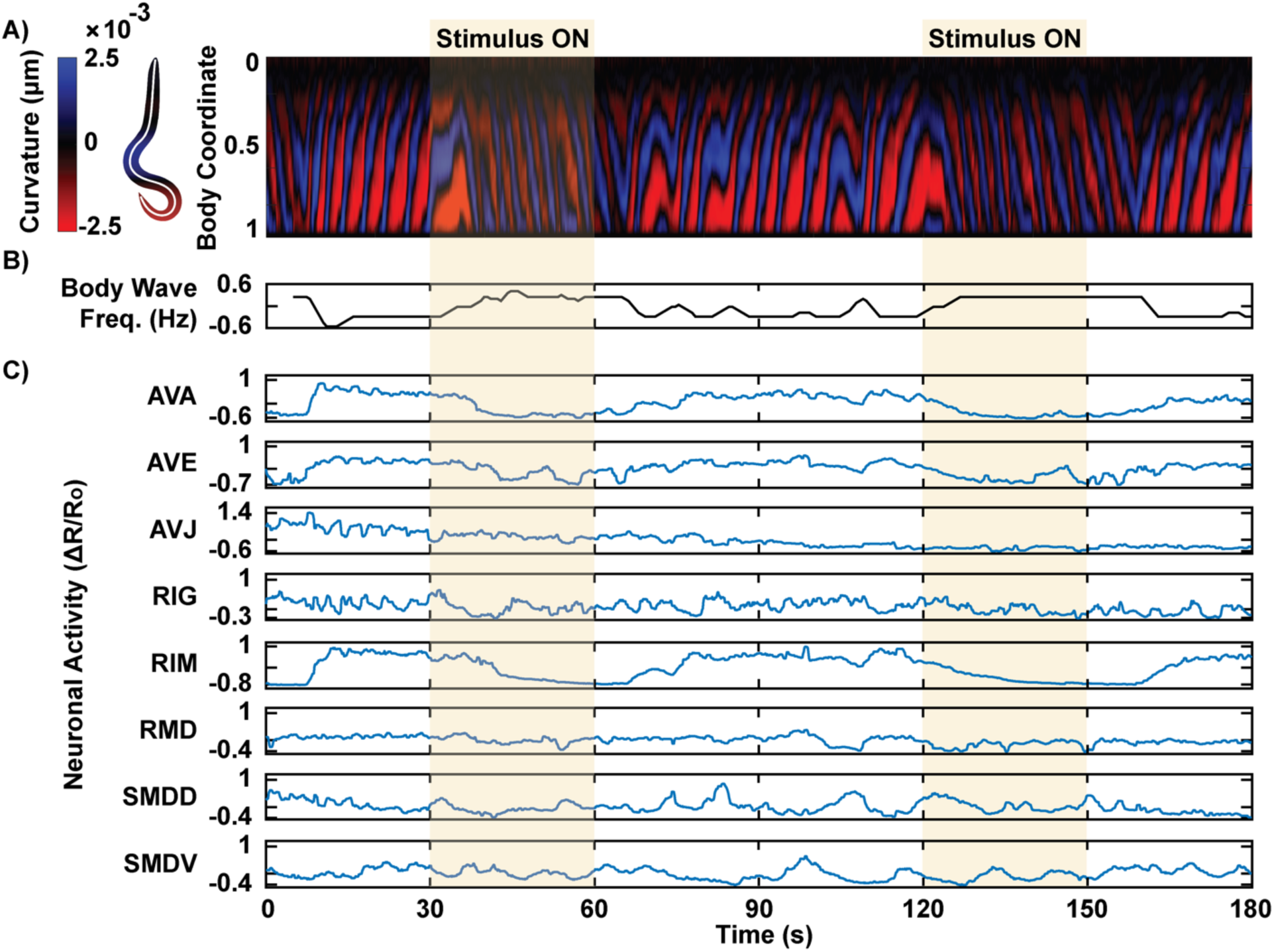
Simultaneous recording of behavior and neuronal activities during chemical stimulation. A) A kymograph of body curvature is generated by segmenting the worm’s centerline. An example recording shows an animal changing its wave propagation direction in response to two trials of food odor stimulus. A wave propagating from head to tail (negative slope of the colored bands) indicates forward movement. B) Signed frequency of the body wave is extracted from the kymograph, with positive values representing forward wave propagation and negative values indicating backward propagation. C) Corresponding neuronal activity recordings show correlations with the observed behavior.

To quantitatively analyze behavior, the signed body wave frequency was extracted from the kymograph using short-time 2D Fourier transform, which computes the sinusoidal frequency and phase within sliding windows of the data over time. This approach is particularly robust against noise in the kymograph, which can disrupt other methods, such as thresholding the curvature for specific body coordinates. Unlike thresholding, the 2D Fourier transform considers the wave more holistically across the entire body, providing a more accurate and noise-resistant measure of wave frequency. A positive body wave frequency indicates the frequency of waves traveling from head to tail, corresponding to forward locomotion, while a negative value indicates backward locomotion (Figure 4B). For food odor stimulation, high body wave frequency correlated with the presence of the odor, suggesting the animal’s attempt to move toward the stimulus.

Using the head-fixed device, calcium activity was recorded simultaneously in a transgenic *C. elegans* strain expressing GCaMP and mCherry driven by a *glr-1* promoter in a group of head neurons of interest ^[53]^. This strain includes interneurons and motor neurons involved in locomotion, allowing us to study sensorimotor integration. Calcium signals were measured from GCaMP fluorescence and normalized by the calcium-insensitive mCherry signal to correct for motion artifacts. Neurons were identified based on their stereotypical positions within the head among a predefined list of candidates for multi-cell expression. Neurons with less confident identities were excluded from the analysis. Notably, interneurons associated with backward movement, such as AVA, AVE, and RIM, exhibited a negative correlation with body wave frequency (Figure 4C). The neurons were identified by their stereotypical positions in the head among a list of candidate neurons for the multi-cell expression. Neurons with less confident neuron identities were omitted in the analysis. Observing behavior and neuronal activity side by side allowed for the identification of specific neurons involved in behavior execution.

### Comparative Analysis of Neuronal Activity in Fully-Restrained vs. Head-Fixed Animals

To determine whether head fixation alters neural activity patterns, we next compared the neuronal activity of fully restrained worms in a conventional device with that of head-fixed worms in our device. To do so, confocal fluorescence microscopy was used for volumetric calcium imaging in a transgenic *C. elegans* strain expressing GCaMP and mCherry in a group of head neurons, comprising a mixture of interneurons and motor neurons involved in locomotion. Because the confocal microscope provides higher spatial resolution and improved cell segmentation at the expense of field of view, behavioral responses were not recorded in either device.

We recorded the neuronal response to an attractive food odor stimulus across multiple animals and trials for both the conventional and head-fixed device (Figure 5A). For many neurons, the responses were consistent between the two setups, with no significant differences observed (Figure 5B). For instance, AVA, AVD, and RIM neurons exhibited a stimulus-correlated decrease in activity that were comparable in both devices. In comparison, AVE and RME showed statistically significant differences in mean activity between the two groups (Figure S2, S3). AVE, a neuron involved in backward locomotion ^[47,54]^, exhibited a more pronounced stimulus-correlated decrease in activity in head-fixed worms, while this response was less prominent in fully-restrained worms. Conversely, RME, a neuron involved in forward locomotion ^[35,47]^, displayed a stimulus-correlated increase in activity only in head-fixed worms. In both neurons, the restricted body movement appeared to result in reduced neuronal activity in response to the stimulus.

**Figure 5:**
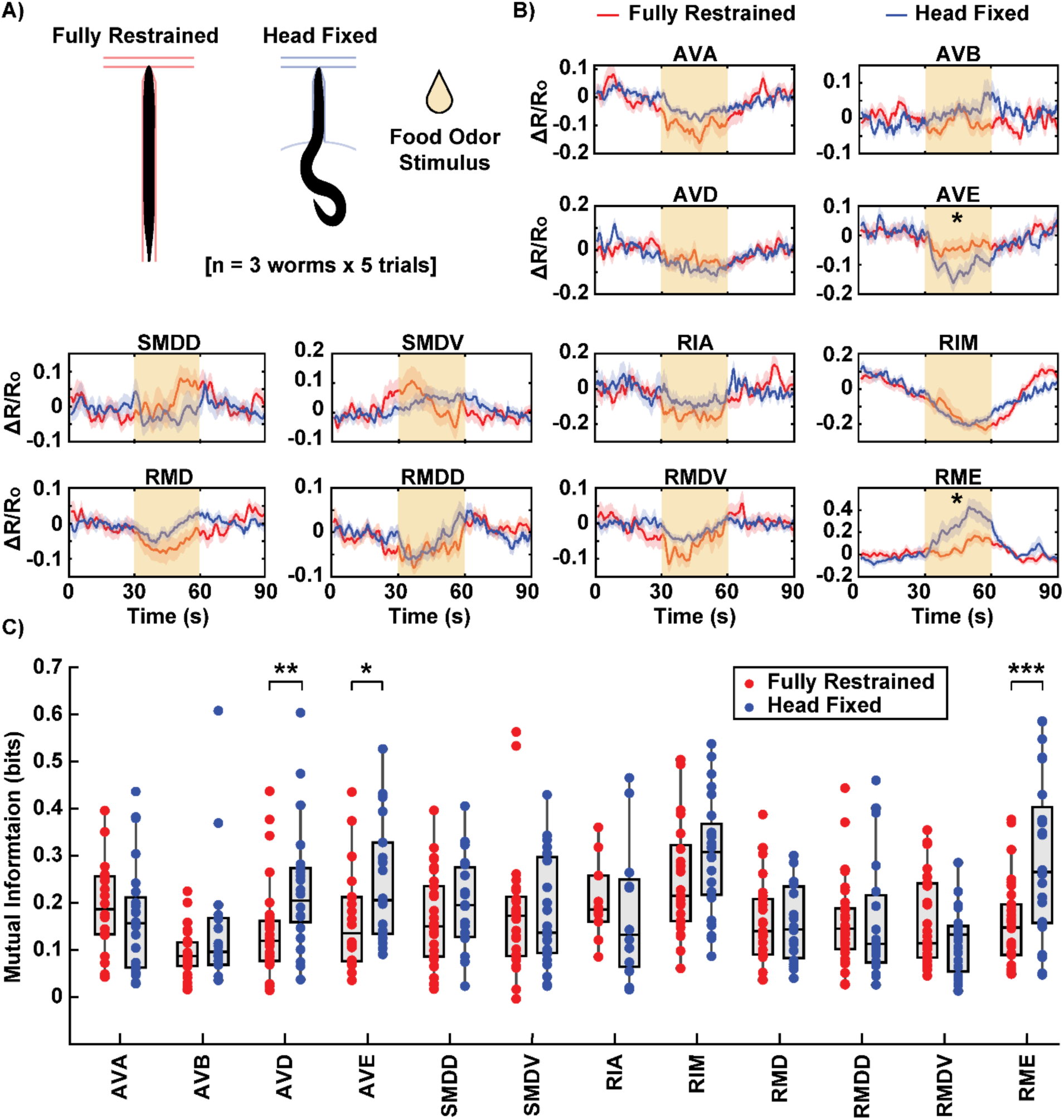
Comparison of neuronal activity in fully-restrained and head-fixed animals. A) A conventional microfluidic device for *C. elegans* chemical stimulation was used to image fully-restrained animals for comparison. Dataset includes three worms per device, with five trials of food odor stimulus per worm. B) Comparison of neuronal activity (ΔR/R0) between fully-restrained (red) and head-fixed (blue) worms. The solid lines represent the average activity over trials for all worms, with shading indicating the standard error of the mean (SEM). The yellow shaded box at 30-60 s indicates food odor stimulation period. Neurons marked with an asterisk (AVE and RME) exhibit a statistically significant difference in mean activity during the stimulus period. C) Mutual information between neuronal activity and the stimulus profile. Each dot represents an individual stimulation trial. The box plot displays the median and the lower and upper quartiles. Certain neurons (AVD, AVE, and RME) show significantly higher mutual information in head-fixed worms compared to fully-restrained worms. Two sample t-tests were performed for statistical analysis. The asterisk symbols denote a significance level as follows: P < 0.05 = *, P < 0.01 = **, P< 0.001= ***.

To better quantify the neuronal responses to the stimulus, we estimated mutual information, a measure of how strongly a neuron’s activity encodes the stimulus. ^[55]^. Unlike simply averaging the activity to a single value for comparison, mutual information considers the entire temporal profile of neuronal activity, capturing more nuanced relationships between the activity and the stimulus. When we computed mutual information between the neuronal activity trace and the discrete stimulus profile for each trial, we found significant differences in the activities of three neurons (AVD, AVE, and RME) between fully-restrained and head-fixed animals (Figure 5C). Unsurprisingly, AVE and RME, which had significantly different mean activity in response to the stimulus between the two groups, also showed significantly different mutual information values. AVD, which did not show strong differences in mean activity, was nonetheless identified by this analysis, demonstrating the sensitivity of the mutual information approach to temporal features of neural activity that are lost through averaging. All three neurons, which are involved in locomotion, exhibited higher mutual information in head-fixed animals compared to fully-restrained ones.

Intriguingly, this finding suggests that the more flexible environment provided by head restraint allows neurons to respond to stimuli in a more naturalistic and dynamic manner. It is possible that the ability to move the tail provides proprioceptive feedback or engages additional neural pathways that enhance the neural response to stimuli. Conversely, restricted body movement in conventional devices may inhibit neuronal activity, potentially leading to muted or unnatural responses in certain neurons. Interestingly, differences in neural responses were observed in only a subset of neurons, such as AVD, AVE, and RME, even though other interneurons are also implicated in similar locomotion functions. This specificity suggests that these neurons may have a stronger proprioceptive component, limiting their full activation or deactivation when the body is fully restrained. This demonstrates the potential of this experimental framework to decouple the causes and effects of behavior, determining whether behavioral context influences neural dynamics or vice versa.

### Simultaneous Monitoring of Neuronal and Behavioral Responses to Aversive Chemical Stimulus

To further demonstrate the capability of our device for simultaneous monitoring of behavior and neuronal activity, we stimulated a population of worms with glycerol, a known noxious chemical ^[30]^. Our primary goal was to validate the device’s ability to elicit expected responses in well-studied neurons, such as RIM, which is known to increase activity during reversals triggered by aversive stimuli. Additionally, we aimed to explore whether the multi-neuronal recordings would reveal interesting trends in less-characterized neurons. Simultaneously capturing both neural and behavioral data during chemical stimulation has traditionally been challenging due to the technical demands of high-resolution neuronal imaging, whole-body behavioral recording, and precise stimulus delivery. Our approach addressed these challenges by using a standard epifluorescence microscope with a 10x objective, enabling most of the worm’s body to remain in the field of view while maintaining sufficient resolution to distinguish individual neurons. The worm’s autofluorescent body, visible in the green channel, allowed simultaneous recording of behavior and neuronal activity without requiring additional complex instrumentation.

As expected, the stimulus-driven responses were clearly observed in both behavior and neuronal activity. During application of the aversive stimulus, body wave frequency significantly decreased and returned to baseline after stimulus removal, indicating increased backward locomotion (Figure 6B,C). This behavioral reversal reflects a stereotyped avoidance response to noxious stimuli. Neuronal activity also showed strong responses to the stimulus (Figure 6D). For example, RIM, an interneuron known to control reversals, exhibited a sharp increase in activity during exposure to the aversive stimulus. This response contrasts with what we observed during attractive odor stimulation, where RIM activity decreased. Additionally, SMDD and SMDV, which are involved in controlling head bending in an anti-correlated manner, displayed opposite trends during the stimulus: SMDD peaked at the onset of the stimulus and ramped down, while SMDV ramped up and peaked at the offset of the stimulus. These findings demonstrate that head-fixed animals can exhibit appropriate neural and behavioral responses to both attractive and aversive chemical stimuli.

**Figure 6:**
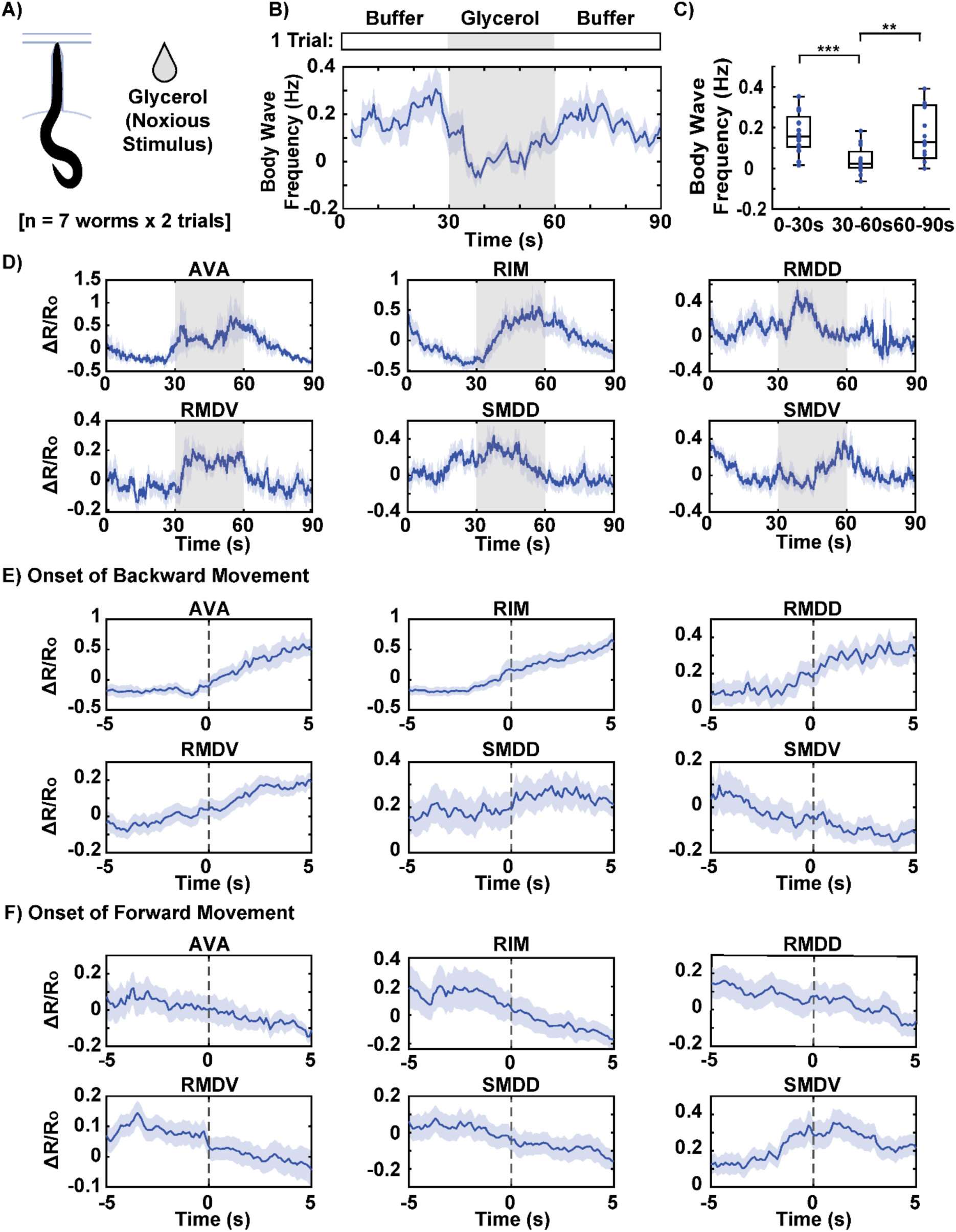
Behavior-coupled neuronal activities in response to noxious stimulus. A) Neuronal activity and behavior of worms in response to glycerol, a noxious stimulus, are recorded simultaneously. The dataset includes 7 worms, with two trials per worm. B, C) The population average of body wave frequency shows a significant decrease during the 30-second glycerol stimulation period, indicating the worms’ aversive response to the stimulus. D) Neuronal traces (ΔR/R0) averaged across worms and trials. E) Average neuronal activity 5 seconds before and 5 seconds after the onset of backward movement. F) Average neuronal activity 5 seconds before and 5 seconds after onset of forward movement. The solid lines represent the average activity, with shading indicating the standard error of the mean (SEM). Two sample t-tests were performed for statistical analysis. The asterisk symbols denote a significance level as follows: P < 0.05 = *, P < 0.01 = **, P< 0.001= ***.

The availability of concurrent behavioral data allows us to interpret neuronal activity within the context of specific behaviors. To begin, we examined the relationship between neuronal activity and behavior across the entire recording, treating all timepoints collectively. By correlating neuronal activity with body wave frequency, we identified neurons that showed either positive or negative correlations with the direction and magnitude of the body’s wave propagation (Figure S4A). For instance, the SMDV neuron displayed a positive correlation with body wave frequency, indicating that its activity increased during forward movement, while other neurons were negatively correlated, suggesting that their activity was associated with backward locomotion. In addition to correlation analysis, we calculated mutual information between neuronal activity and the ternary behavioral states (forward, backward, pause) defined earlier in the study. This analysis allowed us to identify neurons that were more strongly predictive of the behavioral states (Figure S4B). Neurons with higher mutual information values, such as AVA and RIM, exhibited greater predictive power in determining the behavioral state, indicating that these neurons play a key role in encoding and driving distinct motor patterns.

In a more detailed analysis, we examined neuronal activity at the onset of specific behaviors. To do this, we classified worm behavior into forward, backward, and pause states based on thresholding body wave frequency and curvature. In addition to body wave frequency, where the sign clearly indicates forward and backward wave propagation, body curvature was used to identify curling motions. These motions, which exhibit near-zero body wave frequency but represent an important avoidance behavior, were grouped with backward movements, as they are comparable to omega turns in freely moving worms. We aligned neuronal activity to the onset of forward and backward movements, plotting the averaged neuronal activity 5 seconds before and 5 seconds after each transition (Figure 6E,F). For many neurons, activity showed opposing trends during these transitions. For example, AVA and RIM neuron activities increased at the onset of backward movement and decreased at the onset of forward movement. Interestingly, while many neurons showed sustained changes (continuing to increase or decrease after the transition), SMDV showed a transient increase only at the onset of forward movement, a unique pattern that suggests its role in initiating forward motion. This finding is consistent with a previous study, which reported that SMDV activity increases during transitions from reverse to forward crawling, driven by ventrally initiated head bends ^[13]^. These observations underscore the significance of simultaneously observing behavior and neural activity, which allows us to study context-specific neural responses to stimuli and their direct links to observable behaviors. This approach not only validates established roles of certain neurons but also uncovers less well-known dynamics, such as the transient response of SMDV, highlighting the device’s capacity to link neural activity with behavior.

### Simultaneous Whole-Brain Volumetric Imaging and Behavioral Recording

While standard epifluorescence microscopy is sufficient for simultaneously observing behavior and multi-neuronal activity using our device, the advantages of the device are most fully leveraged when combined with volumetric microscopy and a dual-camera setup. By immobilizing the worm’s head, our device enables higher magnification imaging compared to other microfluidic platforms that rely on wide-field views to image freely roaming worms. This configuration allows for multi-neuronal imaging with enhanced clarity and precision. However, single-camera epifluorescence microscopy presents limitations when imaging multi-neuronal activity: maintaining a wide field of view to capture the tail restricts magnification and reduces fluorescence signal quality, while integrating signals from all z-planes prevents resolving neurons that overlap in depth. This is acceptable for sparsely labeled strains but problematic for dense, whole-brain imaging. To overcome these constraints, we paired the device with a light sheet microscope equipped with a dual-camera system ^[56]^. This setup eliminates the compromise between high-resolution neural imaging and behavioral recording. Using a 40x objective, the head region is imaged volumetrically, while a separate IR camera records behavior at lower magnification. In addition, light sheet microscopy offers several advantages, including fast temporal resolution (20 volumes per second) and reduced photobleaching, as it illuminates only a thin sheet of light. For compatibility with the light sheet microscope, the microfluidic device was bonded to fluorinated ethylene propylene (FEP), which has a refractive index close to that of water, ensuring optimal imaging quality.

To demonstrate the synergistic capability of the device and light sheet microscopy, we recorded whole-brain neural activity in head-fixed *C. elegans* during presentation of a food odor stimulus (Figure 7A). Neurons across the head region were volumetrically imaged using a pan-neuronal GCaMP reporter (Figure 7B). Volumetric imaging resolves neurons that overlap in the z-direction, expanding the number of distinguishable neurons compared to epifluorescence imaging. In a 90 s recording with stimulation from 30–60 s, activity from 110 neurons was simultaneously captured (Figure 7D). Concurrently, the worm’s behavior was captured with the IR camera, revealing head-to-tail body waves shortly after stimulus onset and a reversal upon stimulus removal (Figure 7C,D). Principal component analysis (PCA) of the population activity showed that the first component (PC1), accounting for 43% of variance, closely followed the body-wave frequency profile, indicating that the dominant population-level activity axis reflects motor state (Figure 7E). This is consistent with prior whole-brain imaging studies demonstrating that *C. elegans* neural dynamics are organized along an intrinsic manifold representing behavioral progression ^[57]^. When projected onto the two-dimensional PC space, the whole-brain activity traced a structured trajectory in which positive body wave frequency, indicative of forward movement, aligned along the positive PC1 axis (Figure 7F). At the single-neuron level, correlations with body-wave frequency were broadly distributed (Figure 7G). Some neurons showed strong positive correlations with locomotion, whereas others were negatively correlated, suggesting complementary or antagonistic contributions of individual neurons to motor encoding (Figure 7H). Together, these findings demonstrate that whole-brain activity in *C. elegans* is globally coordinated and dominated by the locomotor state. The ability to capture both global neural dynamics and behavioral output in real time provides a comprehensive view of how distributed neuronal populations encode motor behavior. This proof-of-concept experiment establishes that our platform enables high-speed whole-brain volumetric imaging during chemical stimulation, fully realizing its potential for integrative neurobehavioral studies.

**Figure 7:**
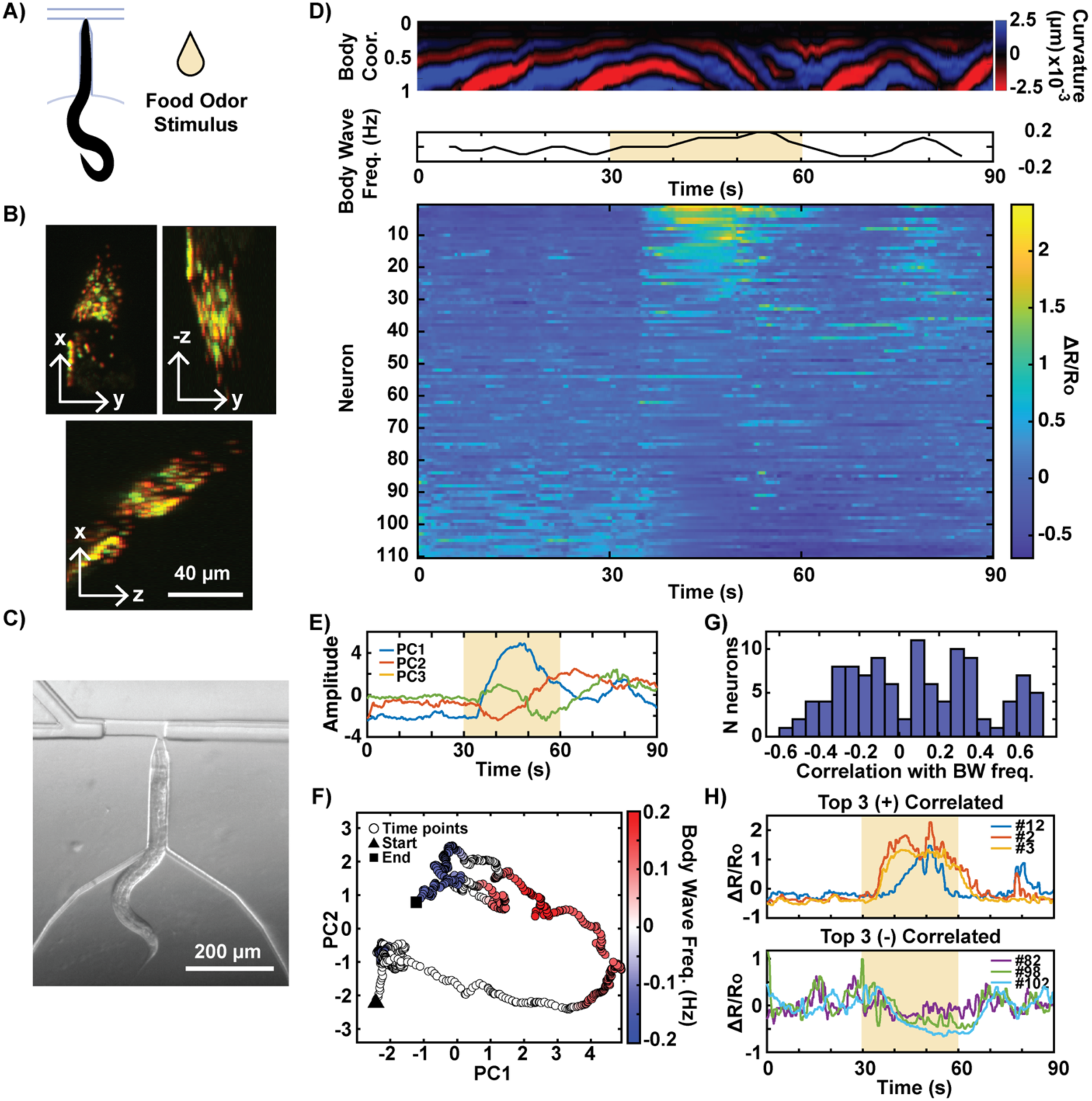
Whole-brain volumetric imaging of neuron activity with simultaneous behavior recording. Light-sheet microscopy enables simultaneous volumetric imaging and behavioral recording within the device. A) A worm’s response to food odor stimulation is shown. B) Volumetric recording of head neurons enables resolution of individual neurons closely clustered or overlapped in the z direction. C) Simultaneous recording of the worm’s behavior at a lower magnification using a separate IR camera. D) Correlation between the worm’s behavior and neuronal activity (ordered by PC1 score). E) Time courses of the first three principal components (PC1–PC3) extracted from the whole-brain calcium imaging traces. F) Neural population trajectory projected onto PC1–PC2 space and colored by body-wave frequency. G) Distribution of Pearson correlation coefficients between individual neuronal activity and body-wave frequency. H) Traces of top three neurons most positively and negatively correlated with body-wave frequency (indices in panel D).

## Discussion

In this paper, we developed a novel microfluidic platform that enables simultaneous observation of neural activity and behavior in *C. elegans* during chemical stimulation. To our knowledge, this is the first device to combine chemical stimulation, multi-neuronal imaging, and behavior recording. By selectively immobilizing the worm’s head while allowing its tail to move freely, we provide a unique system for studying the neural basis of how sensory information is processed and translated into behavior. The freedom of movement in the tail facilitated naturalistic behaviors such as forward and backward locomotion, enabling us to correlate neuronal activity with specific motor patterns. Our results show that head immobilization while allowing free tail movement provides enhanced neuronal responsiveness to stimuli compared to fully immobilized setups. This finding suggests that behavioral context significantly impacts neural circuit function, and the conventional method of chemical stimulation in *C. elegans*, in which the whole body is restrained, may provide unnatural activities in certain neurons, especially those involved in motor control.

While our device addresses many limitations of existing approaches, one inherent caveat is that the animal remains partially unrestrained. The head-immobilized configuration, although necessary for stable high-resolution imaging and chemical stimulation, does not fully replicate the worm’s natural environment and may introduce unnatural proprioceptive feedback that could influence both neuronal and behavioral responses. To minimize these effects, we implemented both experimental and analytical controls. Experimentally, animals were allowed to acclimate to the device prior to imaging to avoid transient loading-induced responses, a standard consideration in *C. elegans* microfluidic imaging. Analytically, stimulus-evoked neuronal and behavioral responses were evaluated relative to baseline (non-stimulated) activity to reduce bias introduced by potential adaptation to the immobilization. Despite these precautions, no current system allows precise chemical stimulation of freely moving *C. elegans* while simultaneously achieving high-resolution neuronal imaging. Thus, our platform currently represents a practical compromise that uniquely integrates controlled chemical delivery with high-resolution neural and behavioral measurements.

Our findings open several avenues for future research. The platform offers opportunities to extend beyond chemosensation and explore how the *C. elegans* nervous system encodes and integrates other sensory modalities, such as mechanosensation and thermosensation. Although these modalities have been examined in fully immobilized animals, such preparations eliminate behavioral readout and the motor and proprioceptive feedback that are integral to natural sensory processing. By allowing limited body movement while maintaining imaging stability, our platform enables investigation of how sensory circuits generate behavior within a more naturalistic and behaviorally relevant context, where sensory input and motor output are dynamically coupled. In our current whole-brain recordings, individual neurons were not identified due to the inherent difficulty of neuron annotation in large volumetric datasets. Reliable neuron identification in future studies will enable systematic mapping of neuronal subtypes and offer deeper insight into how distinct neurons contribute to chemosensation, motor control, and sensorimotor coordination. Finally, the localized immobilization mechanism based on the Venturi effect demonstrated here could be extended to other model organisms, such as zebrafish larvae or Drosophila, to develop comparable platforms for high-resolution neuronal imaging during naturalistic behaviors.

Taken together, the microfluidic device we developed offers a robust platform for studying the interactions between neural circuits and behavior in *C. elegans*. By allowing chemical stimulation and naturalistic behavior during high-resolution neuronal imaging, our device represents a significant advancement without introducing technical hurdles for adoption of the technique. The improved interpretability of the neuronal and behavioral data gathered with this system is anticipated to advance our understanding of how sensory information is processed and transformed into behavior, with implications for more complex nervous systems.

## Methods

### Device fabrication

The device’s master mold was fabricated using a standard photolithography process as described by San Miguel et al. ^[58]^. The device design features a worm inlet, buffer inlet, two stimulus inlets, and an outlet. The worm imaging channel was 40 µm wide, optimized for imaging day 1 adult animals, with channel lengths of 240, 320, or 400 µm depending on the desired level of body restriction. A negative photoresist (SU-8, MicroChem) was used with dark-field photomasks to create positive relief features across three layers. This multi-layer design was critical for producing a three-step vertical taper at the intersection of the worm imaging channel and the stimulus stream, ensuring a precise fit for the worm’s head or tail, thereby effectively limiting movement in the z-direction. The first (bottom) layer was 16 µm thick, and the second and third layers were each 12 µm, resulting in a total thickness of 40 µm in the non-tapered regions of the master. Additionally, a second master, scaled to 125% of the original size in the x, y, and z dimensions, was fabricated for imaging larger day 3 adult animals.

An on-chip valve was incorporated into the design to better control the loading of a single animal at a time. The valve is opened to transfer a worm from the loading chamber to the imaging chamber and closed to prevent additional worms from entering. To enable microfluidic actuation of the valve, two compositions of polydimethylsiloxane (PDMS, Sylgard 184, Dow Corning) were used, following the protocol provided by Cho et al. ^[21]^. A thin layer of more elastic PDMS (23:1 base to curing agent ratio) was deposited on to the master via spin coating (speed: 1000 rpm, ramp: 5 s, spin time: 30 s), partially cured, and then assembled with a thicker layer of firmer PDMS (10:1 base to curing agent ratio). The assembled layers were cured overnight in an 80°C oven. After curing, the PDMS was cut, hole-punched, and bonded to a glass coverslip using plasma treatment.

### Device operation

The four inlets of the device (Figure 1A) were each connected via tubing to liquid reservoirs, which were connected to a pressure box. Solenoid valves were installed on all four inlet tubes. While large batches of worms can be directly added to the reservoir, the worm inlet tube was fitted with a Luer-lock connector between the device inlet and the reservoir, allowing for loading animals in smaller batches. The pressure settings used during device operation are outlined in the table below (Table 1). In brief, the worm inlet was pressurized to 10–30 psi until a worm entered the device through the loading chamber. The same pressure was used to transfer the worm from the loading chamber to the imaging chamber. Once the worm was in the imaging chamber, both the worm inlet valve and the on-chip valve were closed. The buffer inlet stream, pressurized to 10–15 psi, was then used to orient the worm and guide it into the imaging channel. During imaging, the control and test stimuli were alternately activated and deactivated through precise, timed control of the solenoid valves using a MATLAB script.

**Table 1:** Pressure Settings During Device Operation.

| Operation | Worm inlet | Buffer inlet | Control stimulus | Test stimulus |
| --- | --- | --- | --- | --- |
| **Loading worm** | 10-30 psi | Valve closed | 5 psi | Valve closed |
| **Moving worm to imaging channel** | Valve closed | 10-15 psi | 5 psi | Valve closed |
| **Imaging** | Valve closed | 7-10 psi | 5 psi (valve on/off) | 5 psi (valve on/off) |
| **Removing worm** | Valve closed | <20 psi | 5 psi | Valve closed |

### *C. elegans* strains and maintenance

The *C. elegans* strain used for sparse multi-neuronal imaging is ZC3292 *yxEx1701* [*glr-1p::GCaMP6s*, *glr-1p::NLS-mCherry-NLS*] ^[14]^. The strain expresses cytosolic GCaMP and nuclear mCherry, both driven by the *glr-1* promoter. Labeled neurons include AVA, AVB, AVD, AVE, SMDD, SMDV, RIA, RIM, RMD, RMDD, RMDV, and RME. The transgene is also expressed in several other neurons, but these were omitted from the analysis due to difficulty in accurate neuron identification and/or low expression levels. For whole-brain imaging, the strain used is GT408 *lin-15(n765)* X; *hpls675* [*rgef-1p.:GCaMP6s:: 3xNLS: mNeptune + lin-15(+)l*; *aEx22* [*unc-47p::NLS: CyOFPlegI-13NLS* + *gcy-32p: NLS::CyOFPlegl-13NLS*]. This strain exhibits pan-neuronal expression of GCaMP6s and mNeptune, with additional CyOFP expression in twelve landmark neurons driven by the *unc-47* and *gcy-32* promoters.

### Calcium imaging and data processing

#### - Stimulus preparation

For the attractive odor stimulus, *E. coli* in LB was cultured overnight at 37°C. The measured optical density (OD) ranged from 3.0 to 3.5, and the supernatant from the culture was used as the stimulus. For the aversive stimulus, 0.5M glycerol in S. Basal was used.

#### - Epifluorescence imaging set-up

Imaging data for Figure 5 was recorded using an epifluorescence microscope (Leica DMI3000 B) equipped with a color-selective light engine (Lumencor® SPECTRA X) and a two-channel camera (Hamamatsu ORCA-D2). The light wavelengths for two-color fluorescence imaging and UV illumination were as follows: green (excitation: 485 nm, emission filter: 525 nm), red (excitation: 560 nm, emission filter: 617 nm), and UV (excitation: 390 nm). A 10x objective lens with numerical aperture (NA) of 0.30 was used for all imaging. The imaging rate was approximately 10 frames per second.

#### - Epifluorescence data processing

For the 2D epifluorescence recordings, neurons were detected and tracked semi-automatically using the ImageJ TrackMate plugin. A custom MATLAB script was then used to extract fluorescence intensities for each neuron from the spatially registered red and green channels. For each frame, background fluorescence was subtracted from individual pixel intensities. Fluorescence intensity was calculated as the average of the top 20% of background-normalized pixel intensities within a segmented area defined by a fixed distance from the neuron’s centroid. Neuronal activity was quantified as the normalized green-to-red fluorescence ratio, ΔR/R0, where R represents the green-to-red intensity ratio for a neuron in a single frame, and R0 is the mean green-to-red intensity ratio across all frames.

#### - Volumetric imaging set-up

The imaging data used to compare neuronal activity of animals imaged in the conventional device and our device was recorded using a swept-field confocal microscope (Bruker Opterra II) equipped with an EMCCD camera. A 40x objective lens with an NA of 0.75 was used. The imaging rate was 3.3 volumes per second, with each volume consisting of 10 z-slices spanning the expressed neurons. It is important to note that worm behavior was not recorded due to the high magnification and limited field of view.

Volumetric imaging coupled with behavioral observation was performed using a custom-built light-sheet microscope equipped with dual cameras. For fluorescence imaging, a scientific CMOS (sCMOS) camera (Flash 4.0 v3, Hamamatsu) was used, achieving a frame rate of 20 volumes per second. Each volume consisted of 30 slices, captured using a light sheet generated by a micromirror scanner (ASI Imaging) and an electrically tunable lens (EL-10-30-TC, Optotune). An asymmetric pair of objective lenses were used: a 10x lens with a numerical aperture (NA) of 0.3 for wide-field observation and a 40x lens with an NA of 0.8 for high-resolution imaging (both Nikon). Simultaneously, infrared (IR) recordings of behavior were obtained using a 10x long working distance objective lens (Mitutoyo) with an NA of 0.28 and an NIR CMOS camera (Thorlabs).

#### - Volumetric data processing

For all volumetric recordings, neurons were detected and tracked semi-automatically using Zephir ^[59]^. At least three frames per recording were manually annotated to improve accuracy. The quality of the tracking was manually inspected, and the process was repeated as needed to obtain accurate traces. For the whole-brain volumetric recording, 17 neurons were manually annotated across 10 representative frames, and Zephir was then run to generate initial tracking results. After visual inspection of Zephir’s output across all 900 frames, these partial annotations were refined and designated as ground truth for each frame. Subsequently, the remaining neurons (112 in total) were manually annotated in a single reference frame, and Zephir tracked their positions throughout the entire recording using this frame as the seed and the partial annotations as guide points for predicting cell movement. Quality inspection of the resulting traces identified two failed tracks, which were excluded from further analysis. Green and red fluorescence intensities were extracted using Zephir and then processed with a MATLAB script to compute the normalized green-to-red fluorescence ratio (ΔR/R0).

### Behavior quantification

To qualitatively assess worm behavior over time, each recording was translated into a kymograph of body curvature. Using ImageJ, a binary mask of the worm body was extracted from the green channel recordings, where the body was visible due to autofluorescence. This mask was then skeletonized to approximate the body’s centerline. In MATLAB, the skeleton was fitted to a 50-point smoothing spline, and curvature at each point *n* along the spline was calculated as the reciprocal of the radius (in pixels) of a circle passing through points *n-2*, *n*, and *n+2*.

The temporal frequency of a worm’s movement over time was estimated from each kymograph of body curvature. A 2D Short-Time Fourier Transform (STFT) was applied with a window of 40 frames. The frequency is signed, where a positive value indicates forward wave propagation and a negative value indicates backward wave propagation. The magnitude reflects the frequency of a wave cycle traveling along the body.

### Mutual information

To estimate mutual information between neuronal activity, which is a continuous variable, and stimulation profile or behavior, which are discrete variables, we used a MATLAB script available for analyzing discrete-continuous data relationship ^[55]^. Mutual information quantifies the amount of shared information between two variables, providing a robust metric for assessing the relationship between neuronal activity and stimulus and behavioral states. Unlike traditional approaches that rely on binning, this method uses the statistics of distances between data points and their nearest neighbors to calculate mutual information, preserving the resolution of the data. We used a k-value of 5 for nearest-neighbor estimation, which balances sampling error and coarse-graining, as recommended in the original method.

### COMSOL simulation

A three-dimensional model of the T-junction region of the device (Figure 1A), including the three-step vertical taper, was created to simulate the fluid dynamics. The simulation was conducted under the assumption of laminar incompressible flow of water. Two inlets (worm inlet and stimulus inlet) and one outlet were defined in the model. The inlet pressures were set to 8 atm for the worm inlet and 5 atm for the stimulus inlet, with the outlet pressure fixed at 1 atm. The temperature was maintained at 20°C throughout the simulation to reflect experimental conditions. This setup allowed for the analysis of pressure distribution and flow patterns within the T-junction region, providing insights into the functionality and optimization of the Venturi-based immobilization mechanism.

### Particle velocimetry

To determine the instantaneous flow velocity at any point inside the microfluidic device, we employed a particle image velocimetry (PIV) approach. Calibration beads (SPHERO™ Rainbow), 3.0-3.4 μm in diameter, were mixed into a 1x S. basal solution at a density of approximately 3200 particles/μL. Beads flowing inside the microfluidic device were recorded for 5 seconds at 300 frames per second using a high-speed camera (Phantom® model v4.3) mounted on an inverted microscope (Leica DMI3000 B) with a 10x objective (0.3 NA). The images were pre-processed by subtracting the average image of all frames from each time frame to visualize only the beads. The pre-processed videos were then analyzed using PIVlab, an open-source MATLAB tool for particle image velocimetry, and the resulting velocity vectors were averaged across all frames.

## Acknowledgements

The authors acknowledge the funding support of the U.S. NIH (R01MH130064, R01NS115484, R01AG082039) to HL and the U.S. NSF (1764406) to HL.

## Supporting Information

**Figure S1:**
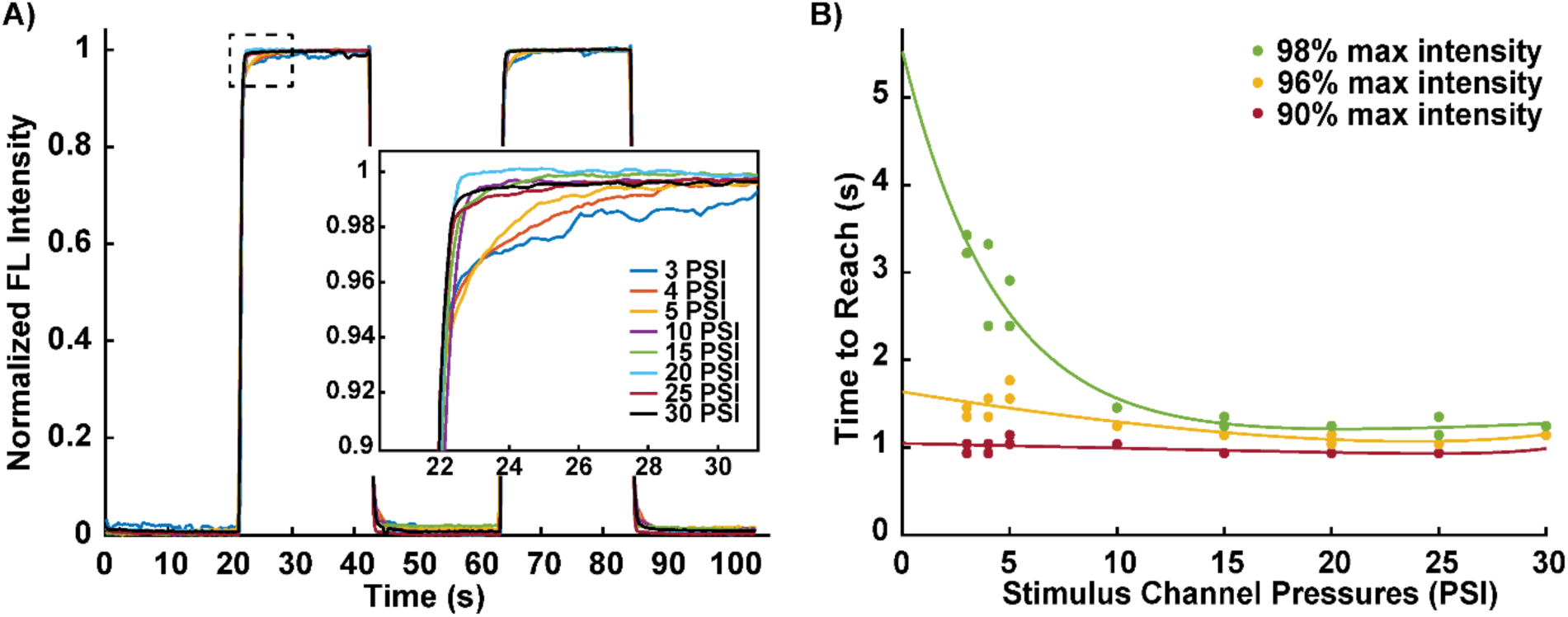
Temporal resolution of chemical stimulation while maintaining head fixation. A) Normalized fluorescence intensity profiles measured near the T-junction during two cycles of fluorescent dye flow at varying stimulus channel pressures. B) Time required to reach 90%, 96%, and 98% of the steady-state (maximum) fluorescence intensity as a function of stimulus channel pressure. Fluid composition switches achieving 90% of steady-state fluorescence or higher can be completed within a few seconds. Solid lines represent exponential fits to the data.

**Figure S2:**
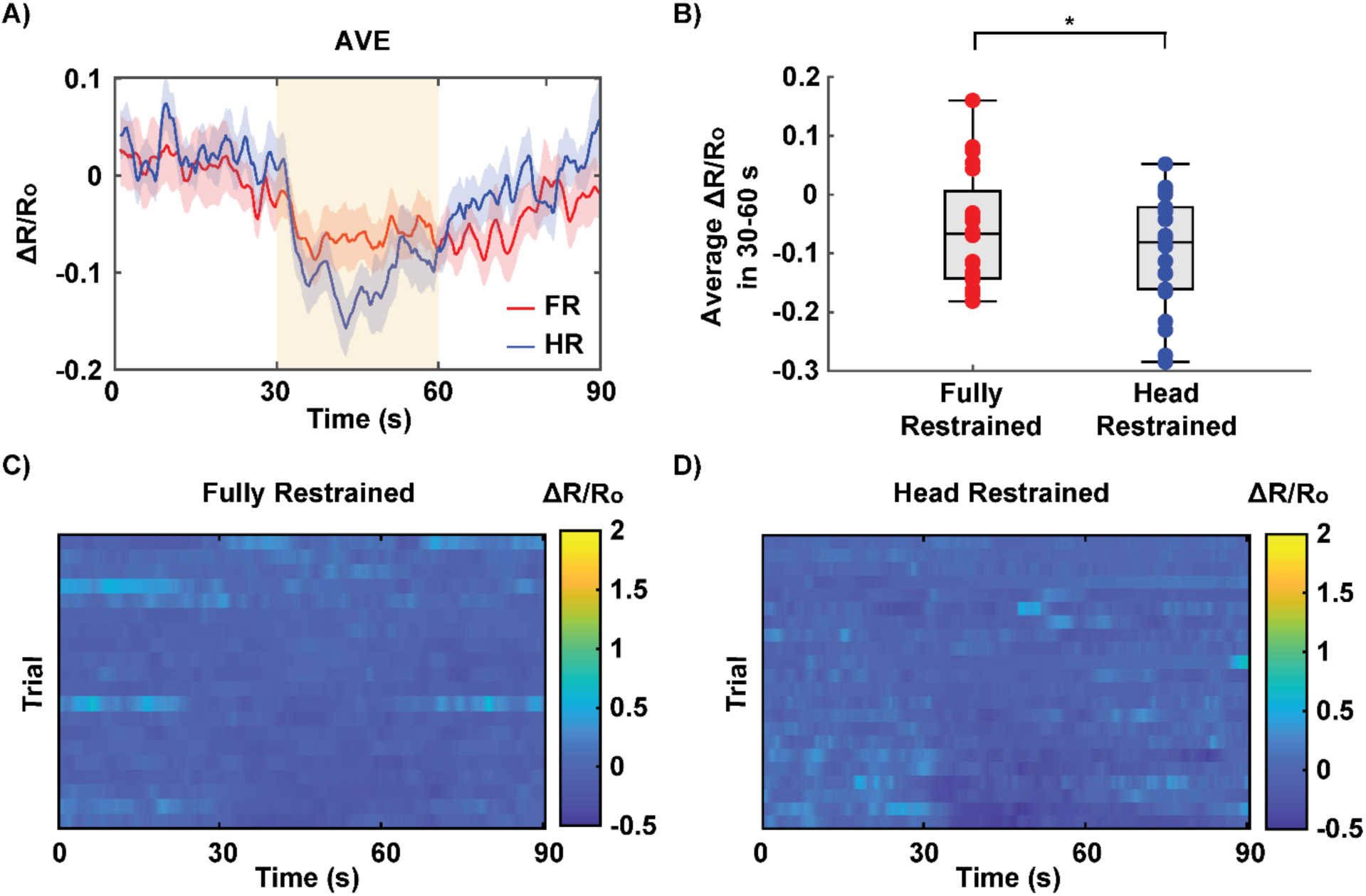
Difference in AVE neuron response in fully-restrained vs head-fixed animals. A) Trial-averaged AVE activity traces (ΔR/R0) of AVE in fully-restrained (FR) and head-fixed (HR) worms. B) Comparison of the average ΔR/R0 values between fully-restrained and head-fixed animals in the 30-60 window. Each dot represents an individual stimulation trial. The box plot displays the median along with the lower and upper quartiles. Statistical significance was determined using two-sample t-tests, with asterisks denoting significance levels: *P < 0.05, **P < 0.01, ***P < 0.001. C,D) Heatmap showing AVE neuron activity (ΔR/R0) across individual trials for fully restrained and head-fixed worms. Each row represents one trial, with the stimulus applied in the 30–60 second window.

**Figure S3:**
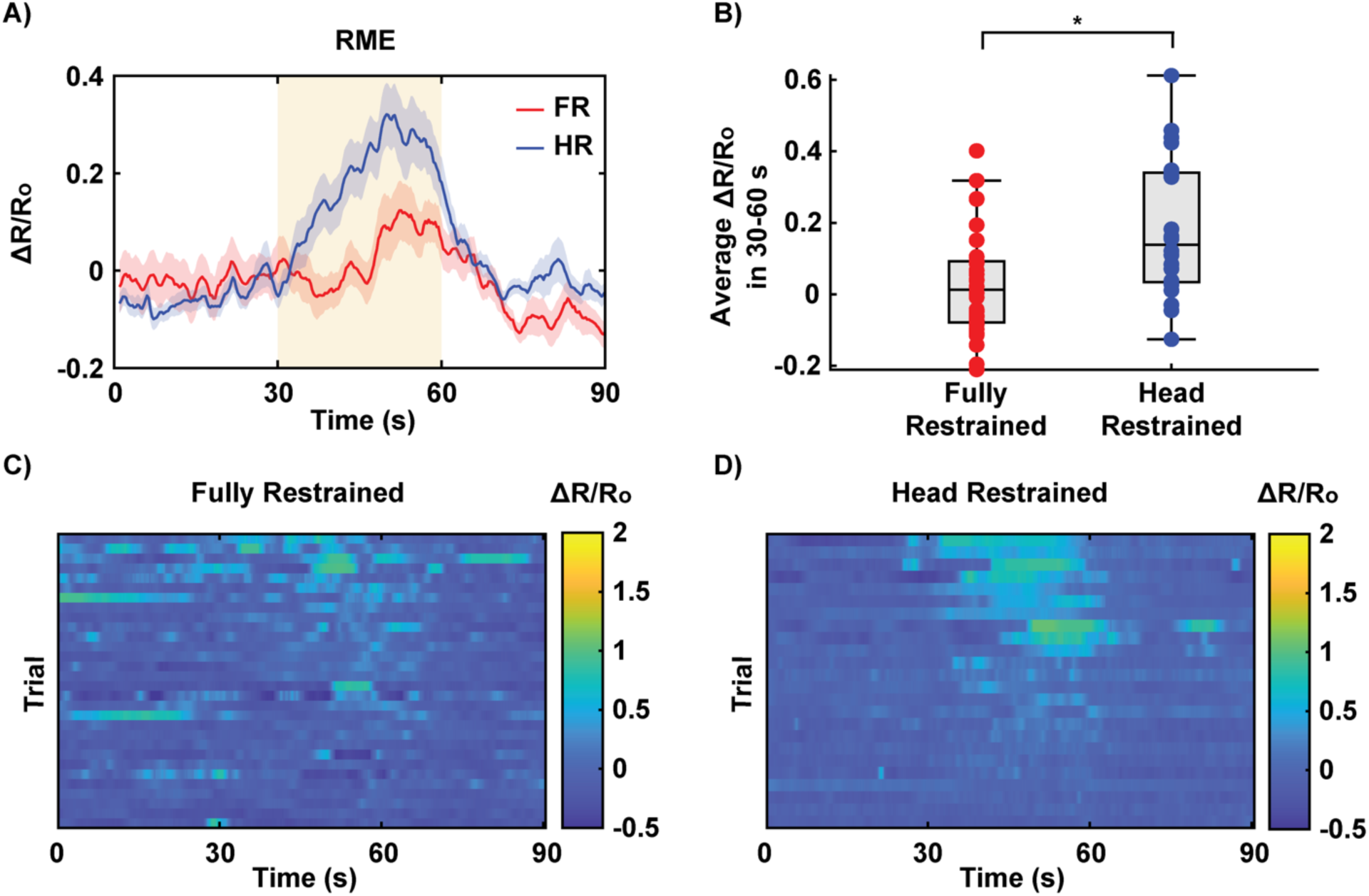
Difference in RME neuron response in fully-restrained vs head-fixed animals. A) Trial-averaged RME activity traces (ΔR/R0) of AVE in fully-restrained (FR) and head-fixed (HR) worms. B) Comparison of the average ΔR/R0 values between fully-restrained and head-fixed animals in the 30-60 window. Each dot represents an individual stimulation trial. The box plot displays the median along with the lower and upper quartiles. Statistical significance was determined using two-sample t-tests, with asterisks denoting significance levels: *P < 0.05, **P < 0.01, ***P < 0.001. C,D) Heatmap showing RME neuron activity (ΔR/R0) across individual trials for fully restrained and head-fixed worms. Each row represents one trial, with the stimulus applied in the 30–60 second window.

**Figure S4:**
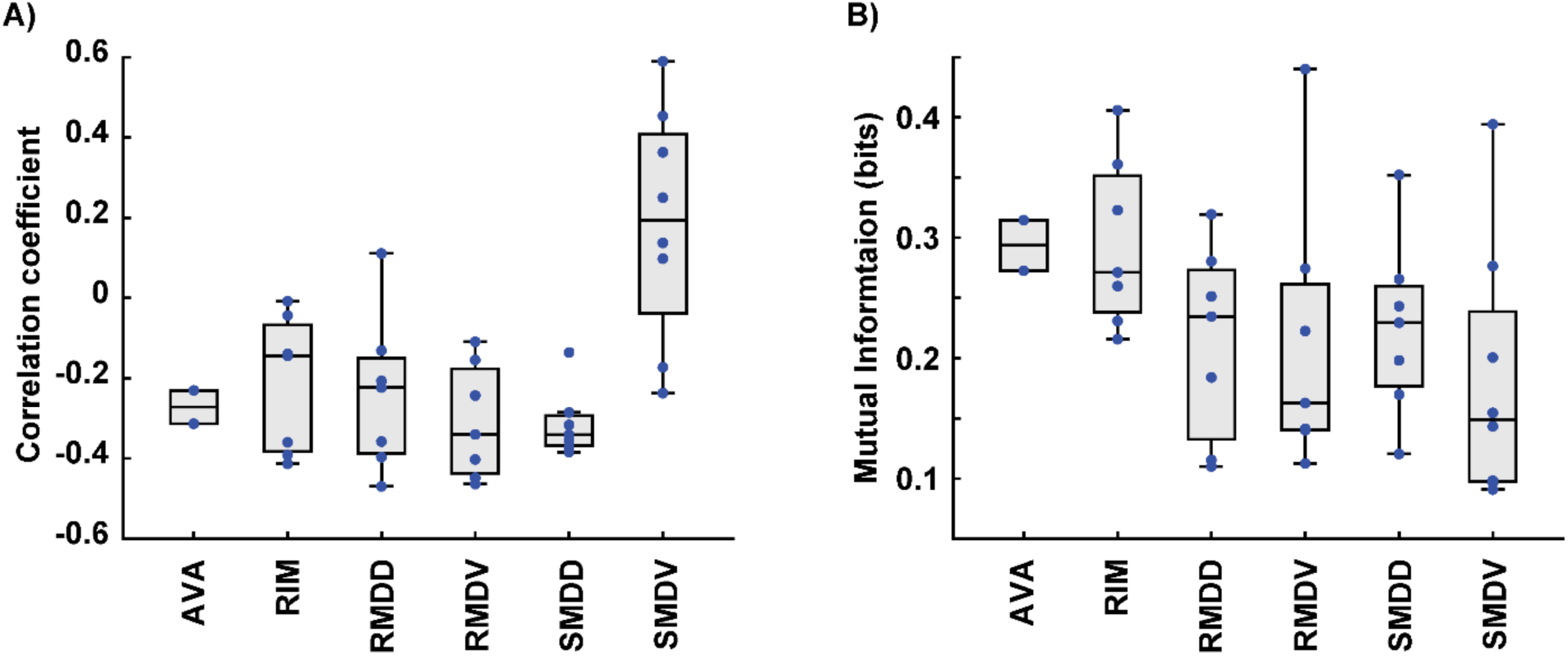
Correlation of neuronal activity with behavior, represented as body wave frequency and ternary classification. A) Pearson correlation coefficients representing the relationship between neuronal activity and body wave frequency for each neuron. Each dot corresponds to a neuron trace from a recording consisting of two trials. A positive correlation coefficient indicates an association with forward movement, while a negative value indicates an association with backward movement. B) Mutual Information between neuronal activity and ternary classification of behavior (forward, backward, and pause). Higher mutual information values indicate a stronger relationship between neuronal activity and the classified behavioral states, showing how much knowing the behavioral states reduces uncertainty about the neuronal activity, and vice versa. The box plots display the median and the lower and upper quartiles.

